# Concerted cellular responses to type I interferon propel memory impairment associated with amyloid β plaques

**DOI:** 10.1101/2021.11.01.466525

**Authors:** Ethan R. Roy, Gabriel Chiu, Sanming Li, Nicholas E. Propson, Hui Zheng, Wei Cao

**Author notes:** Correspondence: Wei Cao One Baylor Plaza, BCM 230, Baylor College of Medicine, Houston, TX 77030, USA.

## Abstract

Despite well-documented maladaptive neuroinflammation in Alzheimer’s disease (AD), the principal signal that drives memory and cognitive impairment remains elusive. Here, we reveal robust, age-dependent cellular reactions to type I interferon (IFN), an innate immune cytokine aberrantly elicited by β amyloid plaques, and examine their role in cognition and neuropathology relevant to AD in a murine amyloidosis model. Long-term blockade of IFN receptor rescued both memory and synaptic deficits, and also resulted in reduced microgliosis, inflammation, and neuritic pathology. Interestingly, microglia-specific IFN receptor ablation attenuated the loss of post-synaptic terminals, whereas IFN signaling in neural cells contributed to pre-synaptic alteration and plaque accumulation. Intriguingly, IFN pathway activation displayed a strong inverse correlation with cognitive performance, promoting selective synapse engulfment by microglia rather than amyloid plaques. Overall, IFN signaling represents a critical module within the neuroinflammatory network of AD and prompts a concerted cellular state that is detrimental to memory and cognition.

## Introduction

Alzheimer’s disease (AD) is the main cause of dementia, characterized by memory impairment. Hallmarked by the deposition of β-amyloid plaques and accumulation of neurofibrillary tangles, AD pathogenesis manifests with complex interactions between different brain cell types (1). Collective histological, bioinformatic and molecular analyses have highlighted a perpetual activation of microglia, the brain resident immune cells, and remarkable connection of a number of AD risk polymorphisms and rare variants to microglia and innate immunity (2, 3). Despite an overwhelming consensus on the importance of neuroinflammatory responses, the core signal that disrupts cognition and memory in AD is not yet well understood.

Recently, we described a prominent antiviral immune response by microglia in multiple murine amyloid β models, as well as human AD (4). At the center of this branch of innate immunity are type I interferons (IFNs) and a large number of IFN-stimulated genes (ISGs), which usually confer an antiviral state in host cells. However, more light is being shed on the functions of these molecules in sterile central nervous system (CNS) inflammation. Recently, Hur et al. reported that IFITM3, an ISG, functions as an immune switch to increase γ -secretase activity, promoting APP cleavage and amyloid pathology (5). Meanwhile, microglial subsets with gene signatures of IFN response (“interferon-responsive microglia,” or IRMs) have been identified from single-cell RNA-seq (scRNA-seq) studies on murine amyloid β models and human AD brains (6, 7). Moreover, polymorphisms in several ISGs were recognized as risk factor for AD (8), while upregulated IFN response was detected in AD patients carrying the *TREM2* R47H variant (9). Given these significant findings, in-depth analysis of the functional contribution of IFN pathway to AD pathogenesis is warranted.

We previously focused on young 5XFAD mice, a widely-studied Aβ model, in which microglia innately responded to amyloid fibrils harboring nucleic acids (NA), activated IFN response pathway, and promoted acute, complement-dependent synapse elimination (4). Here, we examined the accrual of IFN-activated microglia over time and assessed heterogeneity of IFN-responsive microglia. To determine the effects of IFN on cognitive function, plaque pathology, and neuroinflammation, we performed a long-term blockade in older 5XFAD mice with abundant plaques. We further analyzed 5XFAD mice deficient of IFN receptor in different cell lineages to reveal cell-type specific roles of IFN signaling. Overall, we find that IFN signaling via multiple cell types is essential for memory impairment and synaptic damage during amyloidosis.

## Results

### Age-dependent cellular activation by IFN in amyloidosis

Previously, we observed an age-dependent increase of NA-containing plaques in 5XFAD brains (4). Nucleic acids, when complexed to amyloid, serve as an immunogenic stimulus that elicits an IFN response (10). We thus detected microglia with active IFN signaling, marked by nuclear Stat1, exclusively near NA^+^ amyloid plaques in young 5XFAD brains. To comprehensively examine the brain cells activated by IFN, we first generated a reporter mouse line (MxG) by crossing Mx1-Cre mice with the *ROSA26*^mT/mG^ strain, in which IFN exposure results in *Mx1* (a well-known ISG) promoter-driven permanent GFP expression in responsive cells (11). These mice were then bred into the 5XFAD background and examined at different ages to gauge IFN signaling in the brain.

At 3 months, only a small number of GFP^+^ cells were detected, most of which were identified as plaque-associated microglia, making up about 21% of all microglia in plaque-bearing regions (Fig. 1a,b). By 5 months, GFP^+^ microglia became more prevalent and, by 11 months of age, GFP^+^ cells represented a majority of the microglia in these plaque-rich regions. We previously showed that Axl protein, a receptor tyrosine kinase (RTK) and known ISG, is enriched in both Stat1^+^ microglia surrounding amyloid plaques in mice, and neuritic plaques in human brains (4). As expected, we observed significant overlap of GFP and expression of Axl in microglia (Fig. 1c). In β-amyloidosis, a subset of microglia adopt a disease-associated microglia (DAM) phenotype, marked by high Clec7a expression (12, 13). To estimate the heterogeneity of IFN responsiveness within the DAM population, we quantified proportions of microglia in plaque-loaded regions of 5-month-old 5XFAD;MxG brains using GFP and Clec7a expression, and found that roughly half of Clec7a^+^ cells were GFP^+^ (Fig. 1d). This is consistent with elevated ISG transcripts detected in bulk Clec7a^+^ microglia transcriptome from APP-PS1 mice (4), and more importantly reveals a distinct subpopulation within DAM marked by IFN responsiveness. Interestingly, small numbers of astrocytes and blood vessels also expressed GFP in 5- and 11-month-old 5XFAD brains (Fig. S1a,b), suggesting IFN signaling goes beyond microglia amid accumulating CNS amyloidosis. Of note, we found that the MxG reporter yielded no GFP expression in neurons after direct brain administration of IFN, though numerous glial cells and blood vessels turned green (data not shown). Therefore, since neurons were unable to express the Mx1-Cre-dependent GFP readout, we later relied on other methods to detect IFN signaling in neurons.

**Fig. 1:**
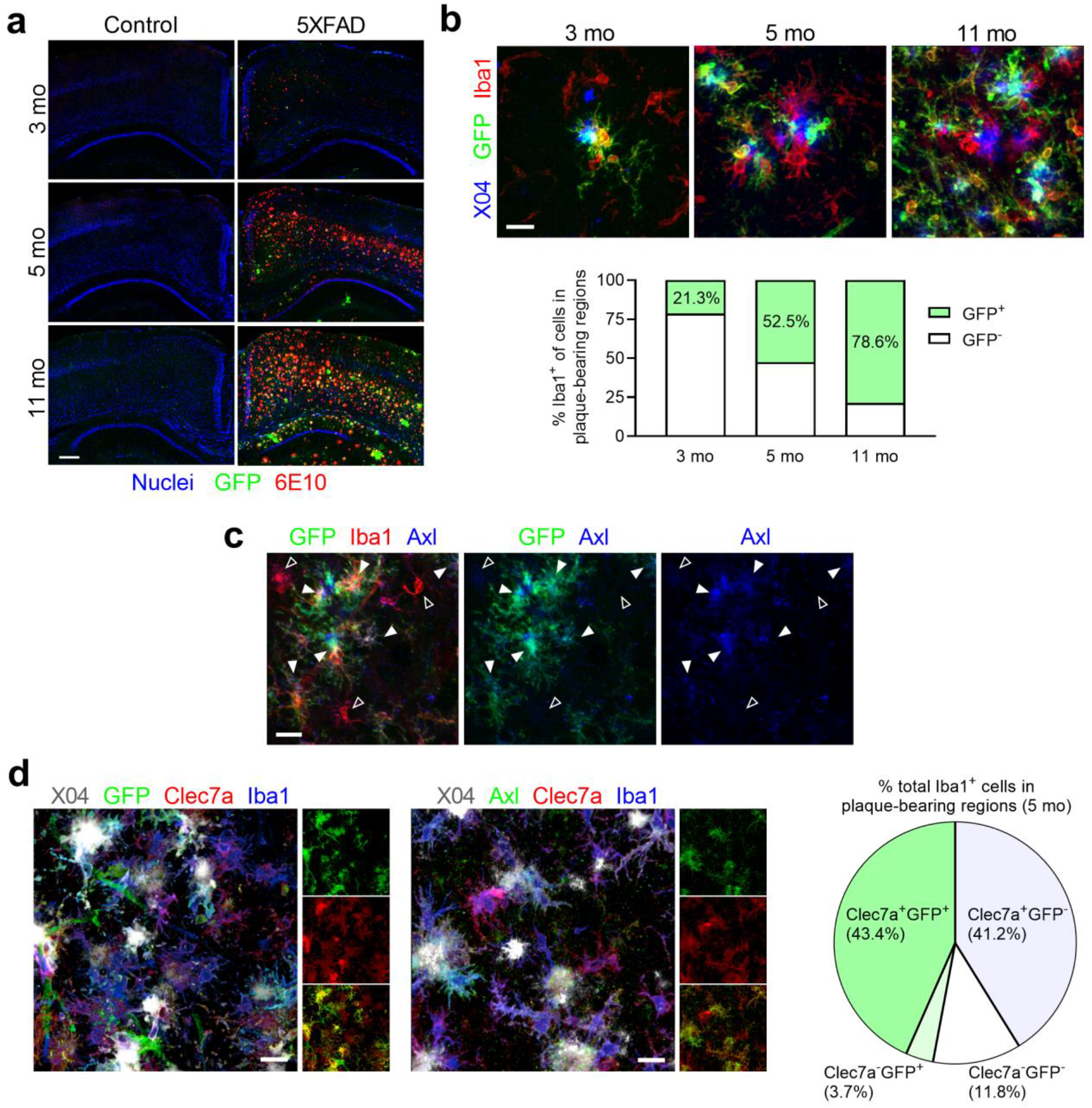
Progressive IFN signaling in 5XFAD brain. **a**, GFP expression upon Mx1-Cre-driven recombination in the mT/mG reporter line crossed to the 5XFAD model. Representative images of 5XFAD and nTg control brains at 3 months (*n* = 3 control, *n* = 3 5XFAD), 5 months (*n* = 3 control, *n* = 3 5XFAD), and 11 months (*n* = 2 control, *n* = 2 5XFAD) of age, showing age-dependent expansion of IFN-responsive GFP^+^ cells throughout plaque-bearing regions (scale bar, 250 µm). **b**, Representative confocal images of tissues from **a** co-labelled with Iba1 with quantification below, showing an IFN-responsive GFP^+^ subset of plaque-associated microglia that expand in an age-dependent manner (3 months: *n* = 75 cells from 3 animals; 5 months: *n* = 301 cells 3 animals; 11 months: *n* = 140 cells from 2 animals; scale bar, 20 µm). **c**, Representative image of a 5-month-old 5XFAD animal (*n* = 3 animals) revealing that Axl is expressed primarily in the GFP^+^ subset of microglia (solid arrowheads) over the GFP^-^ subset (hollow arrowheads). Scale bar, 20 µm. **d**, Representative images from 5 month old 5XFAD brains showing both GFP^+^ and GFP^-^ cells (left) and varying Axl expression (middle) among the plaque-associated Clec7a^+^ subset of microglia, revealing significant heterogeneity of MGnD microglia in the plaque environment (scale bars, 15 µm). (Right) Quantification of microglial subtypes at 5 months using Iba1, Clec7a, and IFN-responsive GFP reporter expression (*n* = 408 cells from 3 animals).

These findings demonstrate a striking age-dependent accrual of IFN-activated brain cells, particularly microglia, and illuminate a degree of heterogeneity among plaque-associated microglia.

### IFN blockade rescues memory and synaptic deficits without altering plaque load

To examine the role of IFN in memory impairment, we implanted osmotic pumps with ventricular cannulae to administer an antibody that specifically blocks signaling of type I IFN receptor (IFNAR) into 4-month-old 5XFAD mice for 30 days (Fig. 2a). Mice were then subjected to Y maze and novel object recognition (NOR) assays to evaluate cognitive aspects related to memory loss, before brain tissues were subjected to detailed histological examination and gene expression analysis. As shown in Figure 2b, 5XFAD mice receiving isotype control IgG failed to show spatial novelty preference in the Y maze, as well as discriminate between novel and familiar objects in the NOR assay, indicating severe deficits in both working memory and short-term reference memory retrieval, respectively. In contrast, 5XFAD mice receiving blocking antibody behaved comparably to non-transgenic control mice, suggesting a full restoration of the memory deficits spurred by amyloid deposition.

**Fig. 2:**
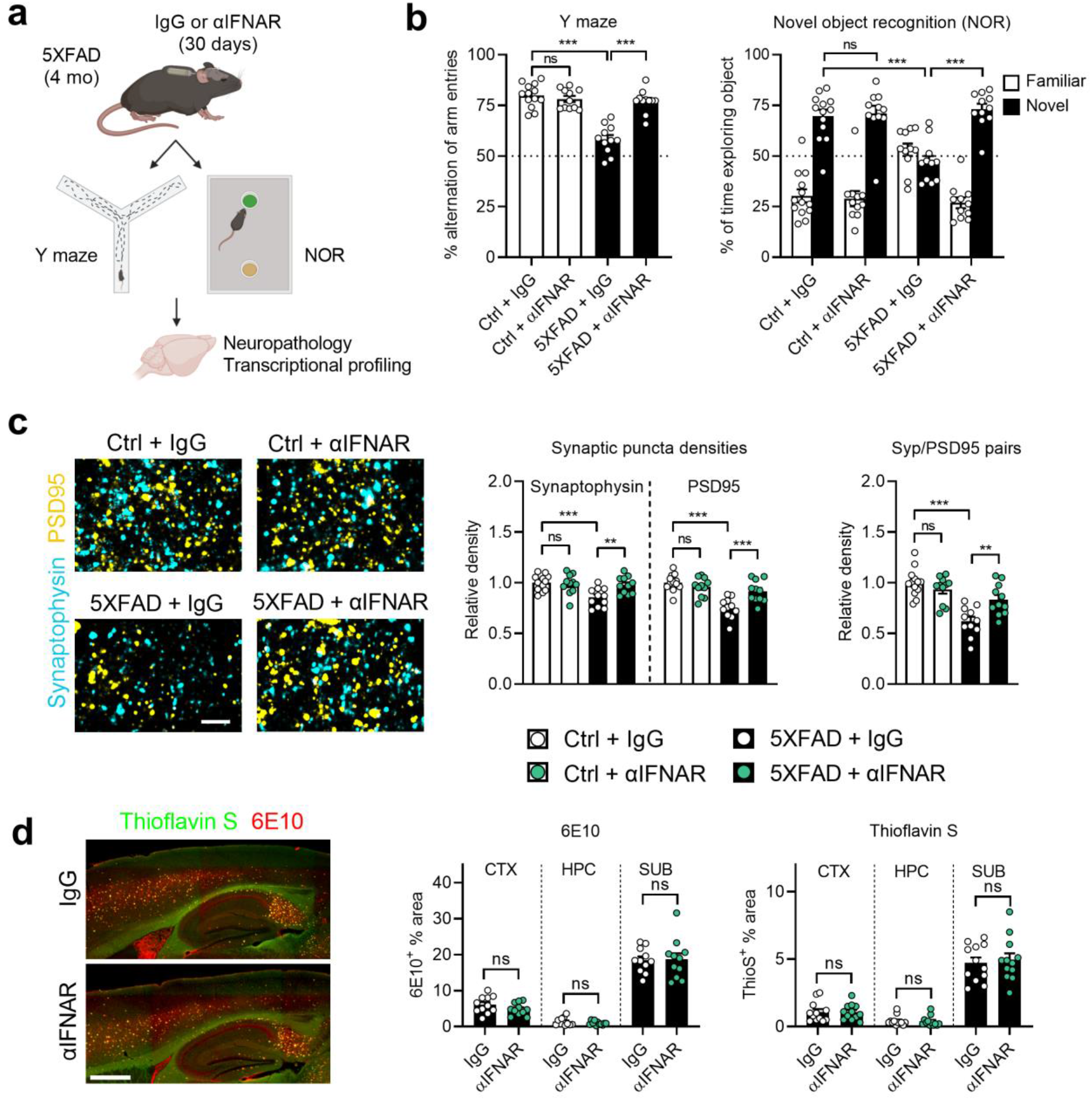
Long-term IFN blockade rescues memory and synaptic deficits without altering plaque load. **a**, Schematic depicting long-term i.c.v. administration of αIFNAR or IgG control to 4-month-old control or 5XFAD mice via mini osmotic pumps with brain ventricular cannulae. After 30 days mice were subjected to behavioral assays to assess memory loss and then harvested for brain tissue analyses. NOR: novel object recognition. **b**, Results of behavioral assays. Ctrl + IgG, *n* = 13 animals; Ctrl + αIFNAR, *n* = 11 animals; 5XFAD + IgG, *n* = 11 animals; 5XFAD + αIFNAR, *n* = 11 animals. Data represent means and s.e.m. Statistics for Y maze were performed with ordinary one-way ANOVA (*P* <0.001, F_42_ = 33.46) and Bonferroni’s multiple-comparisons test. ns, not significant; ^***^*P* < 0.001. Statistics for NOR were performed with two-way ANOVA (*P* < 0.001, F_84_ = 28.36) and Tukey’s multiple-comparisons test. ns, not significant; ^***^*P* < 0.001. **c**, Representative high-magnification images of pre-synapses, marked by synaptophysin, and post-synapses, marked by PSD95, in subicula of treated Ctrl and 5XFAD animals (scale bar, 3 µm). Quantification of relative densities of synaptic markers, and the density of functional synapse pairs (<200nm distance between puncta). Ctrl + IgG, *n* = 13 animals; Ctrl + αIFNAR, *n* = 11 animals; 5XFAD + IgG, *n* = 11 animals; 5XFAD + αIFNAR, *n* = 11 animals. Data represent means and s.e.m. Statistics were performed with ordinary one-way ANOVA (Syp: *P* < 0.001, F_42_ = 6.659; PSD95: *P* < 0.001, F_42_ = 15.36; Co-localized pairs: *P* < 0.001, F_42_= 15.98) and Bonferroni’s multiple-comparisons test. ns, not significant; ^*^\**P* <0.01; ^***^*P* < 0.001. **d**, Histological examination of plaque burden in plaque-bearing regions after long-term administration of αIFNAR using 6E10 antibody to mark Aβ fibrils, and thioflavin S to mark dense core plaques (scale bar, 500 µm). Quantifications of plaque load for both markers in relevant brain regions. 5XFAD + IgG, *n* = 11 animals; 5XFAD + αIFNAR, *n* = 11 animals. Data represent means and s.e.m. Statistics were performed with two-tailed *t*-tests. ns, not significant.

We then performed histological characterization of the brain tissues to explore the neurophysiological basis for the marked reversal of memory loss. Examination of synapse markers showed that 5XFAD mice administered control IgG had reduced levels of synaptophysin and PSD95 proteins, which label pre- and post-synaptic terminals on excitatory neurons, respectively, as well as in functional co-localization (≤200 nm) of these markers (Fig. 2c). Consistent with the outcome of the cognitive assays, these proteins were both significantly elevated in 5XFAD mice with IFN blockade.

To understand the impact of IFN blockade on amyloid pathology, we stained plaques with Thioflavin S and anti-A β antibody but failed to detect a significant difference in dense core plaque load, total A deposition or average plaque volume between 5XFAD mice with and without IFN blockade in any brain region (Fig. 2d; Fig. S2a). Microglia are the primary phagocytes of Aβ species, therefore modifying plaque burden. However, IFN blockade did not affect the amount of Aβ taken up by microglia, nor decrease the level of microglial CD68, a lysosomal receptor involved in phagocytosis (Fig. S2b,c).

Therefore, type I IFN receptor signaling in the brain parenchyma damages memory and synapses during amyloidosis, and suppression of the pathway is sufficient to restore these deficits despite abundant plaque pathology.

### IFN blockade reduces microgliosis, inflammation, and neuritic pathologies

For a deeper understanding of how chronic IFN signaling affects cells in the plaque microenvironment, we examined glia and peri-plaque neuritic structures in detail. IFN blockade effectively reduced Stat1 in the nuclei of both plaque-associated microglia and neurons, indicating broad suppression of excessive IFN signaling in these cells, thus validating the efficacy of the blockade strategy (Fig. 3a). We saw a partial reduction of total Iba1^+^ area with IFN blockade in both cortex and subiculum, the latter an area with the earliest and densest plaque deposition (Fig. 3b). In contrast, astrocyte reactivity markers were not significantly affected in either region (Fig. S3a,b). Further scrutiny of microglia showed that, on a per cell basis, IFN blockade significantly reduced the levels of Axl and Clec7a expression (Fig. 3c), altogether implying an attenuation of microglial activation.

**Fig. 3:**
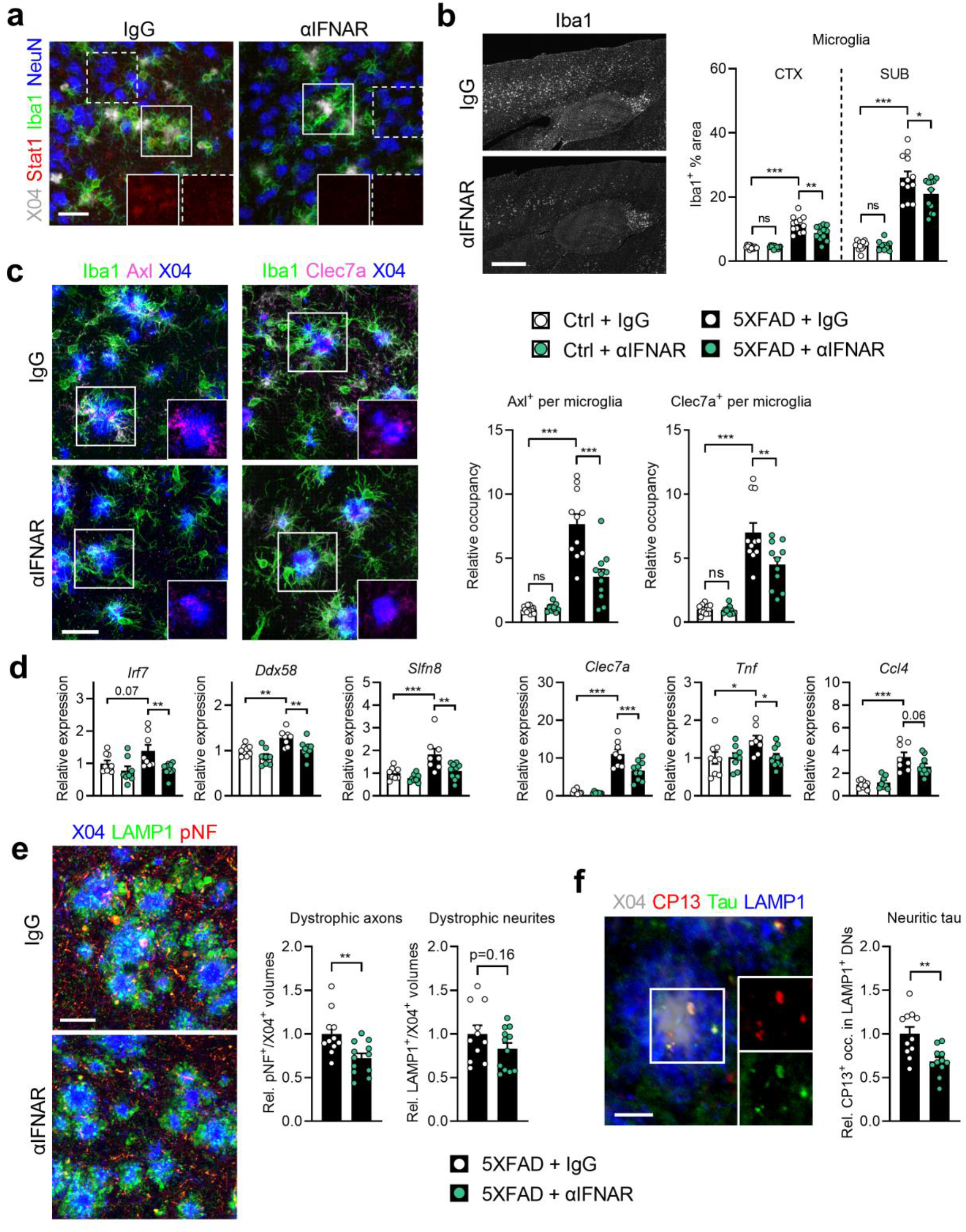
Long-term IFN blockade reduces microgliosis, inflammation, dystrophic axons, and neuritic tau. **a**, Representative images of Stat1 levels, a marker of IFN activation, in the brains of 5XFAD animals treated with IgG (*n* = 11 animals) or αIFNAR (*n* = 11 animals). Insets show isolated Stat1 channels from boxed areas, highlighting microglia (solid boxes) and NeuN^+^ neuronal nuclei (dashed boxes). Scale bar, 30 µm. **b**, Images of Iba1 staining in the cortex and hippocampus of treated 5XFAD animals (scale bar, 500 µm), and quantifications of % Iba1 area. Ctrl + IgG, *n* = 13 animals; Ctrl + αIFNAR, *n* = 11 animals; 5XFAD + IgG, *n* = 12 animals; 5XFAD + αIFNAR, *n* = 12 animals. Data represent means and s.e.m. Statistics were performed with ordinary one-way ANOVA (CTX: *P* <0.001, F_44_ = 50.84; SUB: *P* <0.001, F_44_ = 74.94) and Bonferroni’s multiple-comparisons test. ns, not significant; ^*^*P* < 0.05; ^**^*P* < 0.01; ^***^*P* < 0.001. **c**, Representative images of plaque-associated microglia in treated 5XFAD animals expressing Axl and Clec7a (isolated in insets; scale bar, 30 µm), with quantification of microglial occupancy of both markers. Ctrl + IgG, *n* = 13 animals; Ctrl + αIFNAR, *n* = 11 animals; 5XFAD + IgG, *n* = 11 animals; 5XFAD + αIFNAR, *n* = 11 animals. Data represent means and s.e.m. Statistics were performed with ordinary one-way ANOVA (Axl: *P* <0.001, F_42_ = 42.24; Clec7a: *P* <0.001, F_42_ = 41.77) and Bonferroni’s multiple-comparisons test. ns, not significant; ^**^*P* < 0.01; ^***^*P* < 0.001. **d**, Gene expression alterations with αIFNAR treatment as measured by Nanostring analysis. Ctrl + IgG, *n* = 9 animals; Ctrl + αIFNAR, *n* = 8 animals; 5XFAD + IgG, *n* = 8 animals; 5XFAD + αIFNAR, *n* = 10 animals. Data represent means and s.e.m. Statistics were performed with ordinary one-way ANOVA (*Irf7*: *P* =0.007, F_31_ = 4.923; *Ddx58*: *P* <0.001, F_31_ = 9.283; *Slfn8*: *P* <0.001, F_31_ = 9.086; *Clec7a*: *P* <0.001, F_31_ = 42.99; *Tnf*: *P* =0.036, F_31_ = 3.228; *Ccl4*: *P* <0.001, F_31_ = 18.83) and Bonferroni’s multiple-comparisons test. ns, not significant; ^*^*P* < 0.05; ^**^*P* < 0.01;^***^*P* < 0.001. **e**, Representative images and quantification of LAMP1^+^ dystrophic neurites (DNs) and phospho-neurofilament^+^ (pNF^+^) dystrophic axons (DAs) surrounding methoxy-X04^+^ amyloid plaques in subicula of treated animals (scale bar, 20 µm). 5XFAD + IgG, *n* = 11 animals; 5XFAD + αIFNAR, *n* = 12 animals. Data represent means and s.e.m. Statistics were performed with two-tailed *t*-tests.^**^*P* < 0.01. **f**, Representative image of CP13^+^ aggregated tau foci inside LAMP1^+^ DNs in the subiculum of a 5XFAD animal (insets show isolated channels for CP13 and total tau; scale bar, 30 µm), and quantification of CP13^+^ occupancy in DNs after treatment with IgG or αIFNAR. 5XFAD + IgG, *n* = 11 animals; 5XFAD + αIFNAR, *n* = 12 animals. Data represent means and s.e.m. Statistics were performed with two-tailed *t*-test. ^**^*P* < 0.01.

We next analyzed gene expression in hippocampal tissues from the experimental cohorts and found that IFN blockade not only significantly lowered the expression of ISGs, such as *Irf7, Ddx58* and *Slfn8*, as expected, but also decreased the levels of *Clec7a, Tnf* and *Ccl4*, suggesting a broader dampening effect on neuroinflammation in general (Fig. 3d).

Dystrophic neurites surrounding amyloid plaques represent another hallmark of AD. We found that both phospho-neurofilament^+^ (pNF^+^) dystrophic axons and phosphorylated endogenous tau (CP13^+^ pTau) within dystrophic neurites were partially but significantly diminished by IFN blockade in 5XFAD brain (Fig. 3e,f), implying an IFN-mediated mechanism in promoting these pathologies.

### Selective microglial *Ifnar1* ablation alters microglial activation and prevents post-synaptic loss

All nucleated mammalian cells express the type I IFN receptor and thus can respond to the cytokine. Although our analysis identified microglia as the earliest and primary responder to IFN, how instrumental microglia are in mediating IFN’s overall effects in the brain is not known. We bred *Ifnar1*^fl/fl^ with Cx3cr1*-*Cre^ERT2^ mice, then crossed with the 5XFAD strain to generate mice lacking IFN receptor selectively in microglia (here termed 5XFAD;MKO). When FACS-sorted CD11b^+^ and Cd11b^-^ cells from the brains of MKO mice were analyzed, significant reduction of *Ifnar1* was detected, in conjunction with decreased ISGs, only in the Cd11b^+^ population, confirming the selective knockout in microglia (Fig. S4). Also consistent with this, Stat1 protein was noticeably absent in plaque-associated microglia in 5XFAD;MKO brains (Fig. 4a).

**Fig. 4:**
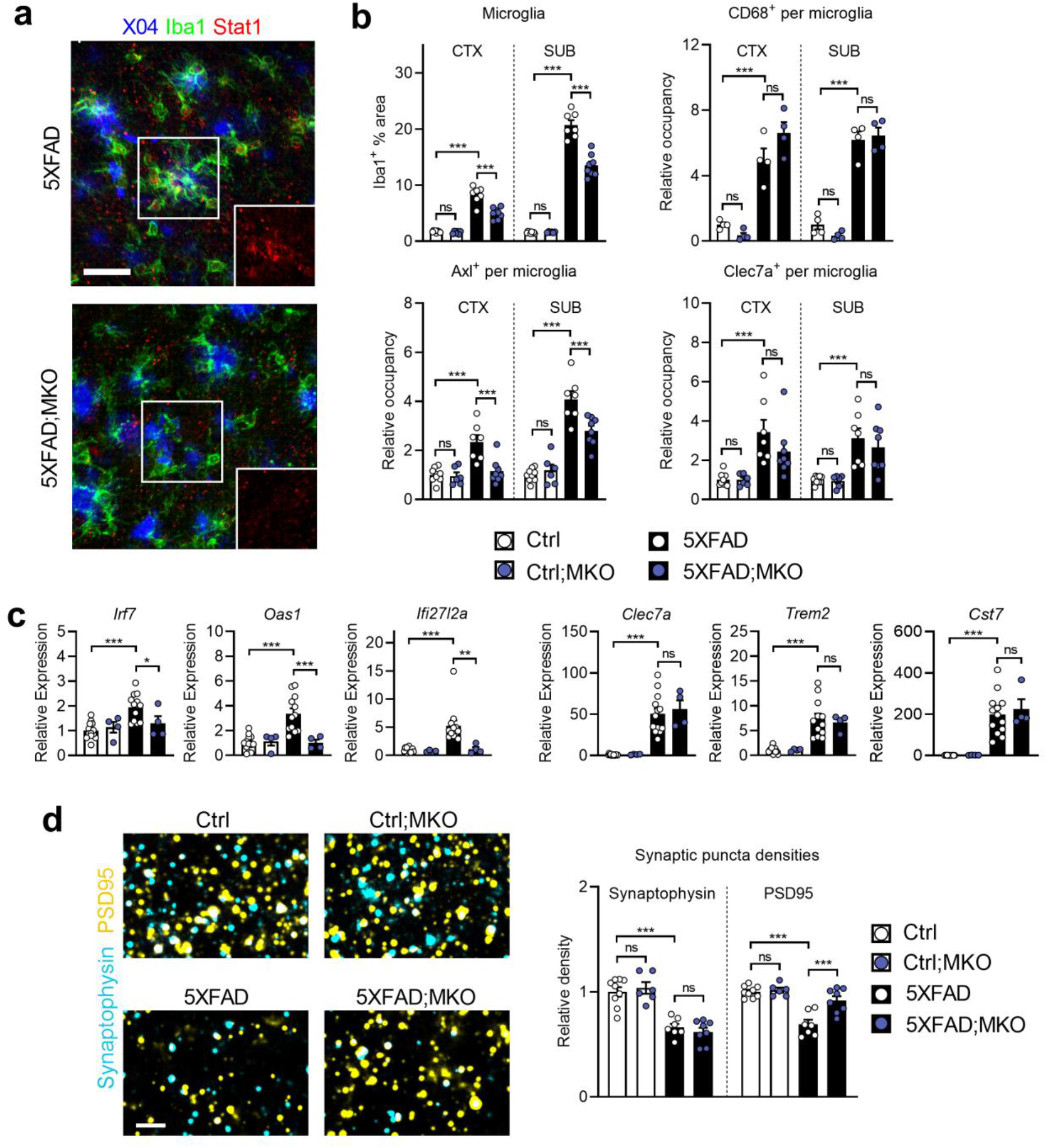
Selective microglial IFNAR ablation alters reactive microglial phenotype and prevents post-synaptic loss. **a**, Representative images of Stat1 expression in the plaque environment of 5XFAD animals with microglia-specific deletion of *Ifnar1* (“5XFAD;MKO”; *n* = 8 animals) compared to *Ifnar1*-sufficient 5XFAD animals (“5XFAD”; *n* = 7 animals) (insets show isolated Stat1 channel in plaque-associated microglia; scale bar, 30 µm). **b**, Quantifications of total % Iba1^+^ area and activation marker occupancy (CD68, Axl, and Clec7a) on a per microglia basis by region. For Iba1, Axl, and Clec7a: Ctrl, *n* = 9 animals; Ctrl;MKO, *n* = 6 animals; 5XFAD, *n* = 7 animals; 5XFAD;MKO, *n* = 8 animals. For Cd68: Ctrl, *n* = 4 animals; Ctrl;MKO, *n* = 4 animals; 5XFAD, *n* = 4 animals; 5XFAD;MKO, *n* = 4 animals. Data represent means and s.e.m. Statistics were performed with ordinary one-way ANOVA (Iba1, CTX: *P* <0.001, F_34_ = 73.93; Iba1, SUB: *P* <0.001, F_34_ = 212.6; Cd68, CTX: *P* <0.001, F_12_ = 38.45; Cd68, SUB: *P* <0.001, F_12_ = 72.61; Axl, CTX: *P* <0.001, F_26_ = 12.52; Axl, SUB: *P* <0.001, F_26_ = 43.68; Clec7a, CTX: *P* <0.001, F_26_ = 8.893; Clec7a, SUB: *P* <0.001, F_26_ = 10.29) and Bonferroni’s multiple-comparisons test. ns, not significant; ^***^*P* < 0.001. **c**, Relative expression of ISGs and microglial activation markers. Ctrl, *n* = 13 animals; Ctrl;MKO, *n* = 4 animals; 5XFAD, *n* = 12 animals; 5XFAD;MKO, *n* = 4 animals. Data represent means and s.e.m. Statistics were performed with ordinary one-way ANOVA (*Irf7*: *P* <0.001, F_29_= 9.300; *Oas1*: *P* <0.001, F_29_ = 13.41; *Ifi27l2a*: *P* <0.001, F_29_ = 11.79; *Clec7a*: *P* <0.001, F_29_ = 27.77; *Trem2*: *P* <0.001, F_29_ = 18.19; *Cst7*: *P* <0.001, F_29_ = 30.97) and Bonferroni’s multiple-comparisons test. ns, not significant; ^*^*P* < 0.05; ^**^*P* < 0.01; ^***^*P* < 0.001. **d**, High-magnification confocal images of pre- and post-synaptic puncta marked by synaptophysin and PSD95, respectively, in subicula of Ctrl (*n* = 9 animals), Ctrl;MKO (*n* = 6 animals), 5XFAD (*n* = 7 animals), and 5XFAD;MKO (*n* = 8 animals). Scale bar, 3 µm. Quantification of relative synaptic puncta densities. Data represent means and s.e.m. Statistics were performed with ordinary one-way ANOVA (Syp: *P* <0.001, F_26_ = 25.05; PSD95: *P* <0.001, F_26_ = 19.92) and Bonferroni’s multiple-comparisons test. ns, not significant; ^***^*P* < 0.001.

Similar to IFN blockade, microglia in 5-month-old 5XFAD;MKO mice showed a reduction of Iba1^+^ area and significantly less Axl expression on a per cell basis (Fig. 4b). However, Clec7a and CD68 levels were not affected. 5XFAD;MKO mice expressed significantly less ISG transcripts, such as *Irf7, Oas1* and *Ifi2712a*, while maintaining the expression of classical DAM markers, such as *Clec7a, Trem2* and *Cst7* (Fig. 4c). These findings suggest that microglial IFN signaling selectively regulates a subset of molecular changes observed in plaque-associated microglia.

Examination of synaptic markers revealed an unexpected effect – while both pre- and post-synapses were reduced in normal 5XFAD brains, PSD95^+^ puncta density, but not synaptophysin^+^, was restored in 5XFAD;MKO (Fig. 4d). We also examined dystrophic neuronal structures and found that pTau levels were significantly reduced by ablating microglia-specific IFN signaling, despite comparable axonal dystrophy (Fig. S5e). Altogether, these results hint at a selective function of IFN signaling in microglial activation and synapse modification.

### Neural *Ifnar1* ablation reduces amyloid plaques and restores pre-synaptic terminals

To understand the importance of IFN signaling in non-microglia cells in the brain, we bred *Ifnar1*^fl/fl^ and Nestin*-*Cre mice to generate 5XFAD mice with the type I IFN receptor ablated in neuroectodermal-derived cells, including neurons and glial cells other than microglia (here termed 5XFAD;NKO). Consistent with the conditional knockout, Stat1 signal in 5XFAD;NKO neuronal nuclei was significantly reduced (Fig. 5a). Of note, we did not detect overt accumulation of LC3B-II or protein hyper-ubiquitination in the brains of adult *Ifnar1*^fl/fl^;Nestin*-*Cre mice as reported (14), nor decreased expression of endogenous or transgenic full-length amyloid precursor protein (APP) with conditional *Ifnar1* ablation, as reported to be affected by germline *Ifnar1* deletion (15) (Fig. S5a).

**Fig. 5:**
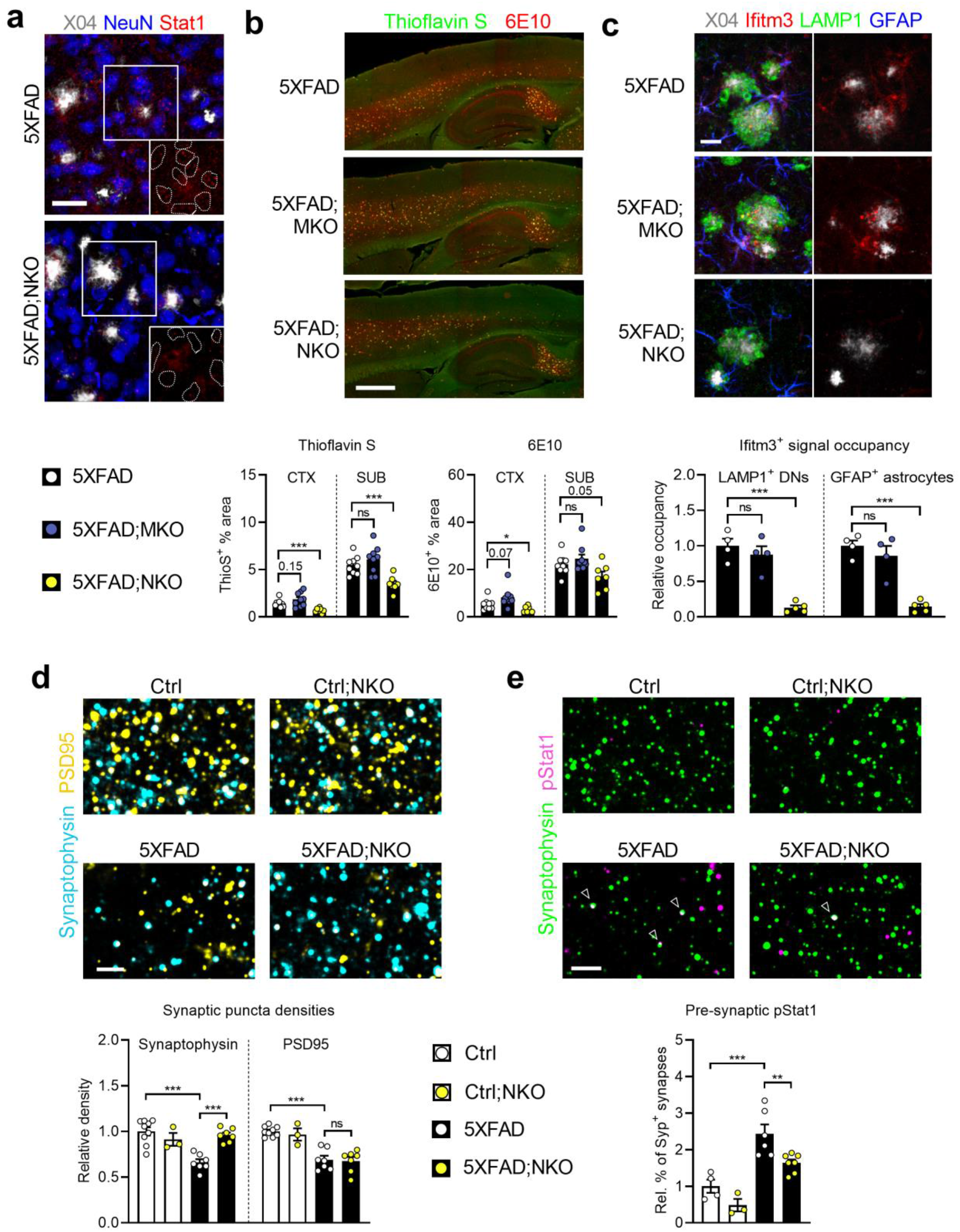
Selective neural IFNAR ablation reduces plaque load and prevents pre-synaptic loss. **a**, Representative images of Stat1 expression in plaque environment in 5XFAD animals with neural-specific deletion of *Ifnar1* (“5XFAD;NKO”; *n* = 7 animals) compared to *Ifnar1*-sufficient 5XFAD animals (“5XFAD”; *n* = 7 animals) (insets show isolated Stat1 channel in outlined NeuN^+^ neuronal nuclei; scale bar, 30 µm). **b**, Histological examination of plaque burden in plaque-bearing regions of 5XFAD (*n* = 9 animals), 5XFAD;MKO (*n*= 9 animals), and 5XFAD;NKO (*n*= 7 animals) using thioflavin S to mark dense core plaques, and 6E10 antibody to mark all Aβ fibrils (scale bar, 500 µm). Quantification (below) of percent area of plaque markers by brain region. Data represent means and s.e.m. Statistics were performed with two-tailed *t*-tests. ns, not significant; ^*^*P* < 0.05; ^**^*P* < 0.01. **c**, Representative images and quantification of Ifitm3 signals localized inside LAMP1^+^ DNs and GFAP^+^ astrocytes (scale bar, 20 µm). 5XFAD, *n* = 4 animals; 5XFAD;MKO, *n* = 4 animals; 5XFAD;NKO, *n* = 5 animals. Data represent means and s.e.m. Statistics were performed with two-tailed *t*-tests. ns, not significant; ^***^*P* < 0.001. **d**, High-magnification confocal images (scale bar, 3 µm) and relative density quantifications of pre- and post-synaptic puncta marked by synaptophysin and PSD95, respectively, in subicula of Ctrl (*n*= 9 animals), Ctrl;NKO (*n*= 3 animals), 5XFAD (*n*= 7 animals), and 5XFAD;NKO (*n*= 7 animals). Data represent means and s.e.m. Statistics were performed with ordinary one-way ANOVA (Syp: *P* <0.001, F_22_ = 14.72; PSD95: *P* <0.001, F_22_ = 21.26) and Bonferroni’s multiple-comparisons test. ns, not significant; ^***^*P* < 0.001. **e**, High-magnification confocal images (scale bar, 3 µm) and quantification of pStat1^+^ puncta co-localized with Syp^+^ synapses in subiculum. Ctrl, *n*= 4 animals; Ctrl;NKO, *n*= 3 animals; 5XFAD, *n*= 6 animals; 5XFAD;NKO, *n*= 7 animals. Data represent means and s.e.m. Statistics were performed with ordinary one-way ANOVA (*P* <0.001, F_16_ = 16.99) and Bonferroni’s multiple-comparisons test. ns, not significant; ^**^*P* < 0.01; ^***^*P* < 0.001.

In contrast to IFN blockade and microglia-specific *Ifnar1* ablation, 5XFAD;NKO mice at 5 months accumulated fewer ThioS^+^ and 6E10^+^ plaques in different brain regions (Fig. 5b). Measurement of A β inside microglial CD68^+^ vesicles indicated unaltered plaque phagocytosis by microglia (Fig. S5b). Since IFITM3 was shown to function as an inflammation-triggered switch to enhance A β production (5), we investigated the possible involvement of this ISG. First, we confirmed a sensitive and dose-dependent induction of Ifitm3 protein in primary neurons by IFN β (Fig. S5c). Further, we confirmed the upregulation of Ifitm3 protein in dystrophic neurites, a known site of heightened Aβ production and release, as well as in astrocytes in 5XFAD brains (Fig. S5d). Interestingly, a selective diminution of Ifitm3 was detected in 5XFAD;NKO, but not 5XFAD;MKO, mice (Fig. 5c). Neural IFN signaling did not have a major impact on overall dystrophic neuronal structures (Fig. S5e).

On synaptic regulation, 5XFAD;NKO displayed an opposing phenotype to 5XFAD;MKO, such that synaptophysin^+^ puncta levels, but not PSD95^+^, were restored, implying a neural-intrinsic and IFN-dependent regulation of pre-synaptic bouton density during disease (Fig. 5d). Activity-dependent events shape neuronal networks in part by elimination of inactive synapses, a mechanism critical for proper configuration of circuits. In post-natal brain, Stat1 signaling at inactive pre-synaptic terminals is instrumental for synapse refinement (16). To gauge the relevance of this axis in pre-synaptic loss during β amyloidosis, we employed an antibody against Stat1 phosphorylated at Tyr701 (pStat1) and detected enhanced frequency of pStat1^+^ pre-synaptic boutons in 5XFAD brain (Fig. S5f,g), suggesting a potential functional involvement. In 5XFAD;NKO subicula, a significantly lower percentage of Syp^+^ pre-synapses were pStat1 positive compared to 5XFAD (Fig. 5e). In accordance, fewer pStat1^+^ puncta were present in the nuclei of CA1 neurons, which project to the subiculum (Fig. S5h).

Overall, these findings reveal pathogenic effects of type I IFN signaling in non-microglial brain cells on plaque formation and synaptic pathology.

### Interrogation of AD-related pathological processes in cognitive performance

Although we detected cell-type specific effects of IFN signaling on multiple AD-related pathologies, the relative importance of each process in the clinically relevant disease manifestation, *i*.*e*. cognition and memory impairment, remains unclear. We thus constructed a database containing gene expression profiles, numerous neuropathological parameters, and behavioral outcomes from the cohort of 5XFAD mice that underwent IFN blockade treatment to perform unbiased correlation analysis. As shown in Figure 6a, ordering profiled genes from strongest negative to strongest positive correlates with performance in Y maze revealed that ISGs were heavily enriched among genes most strongly associated with poor cognition. IRF7 is a master transcriptional regulator of type I IFN-dependent immune response (4). *Irf7* levels were negatively correlated with Y maze performance, highlighting a pathogenic effect of IFN pathway on memory (Fig. 6a,e).

**Fig. 6:**
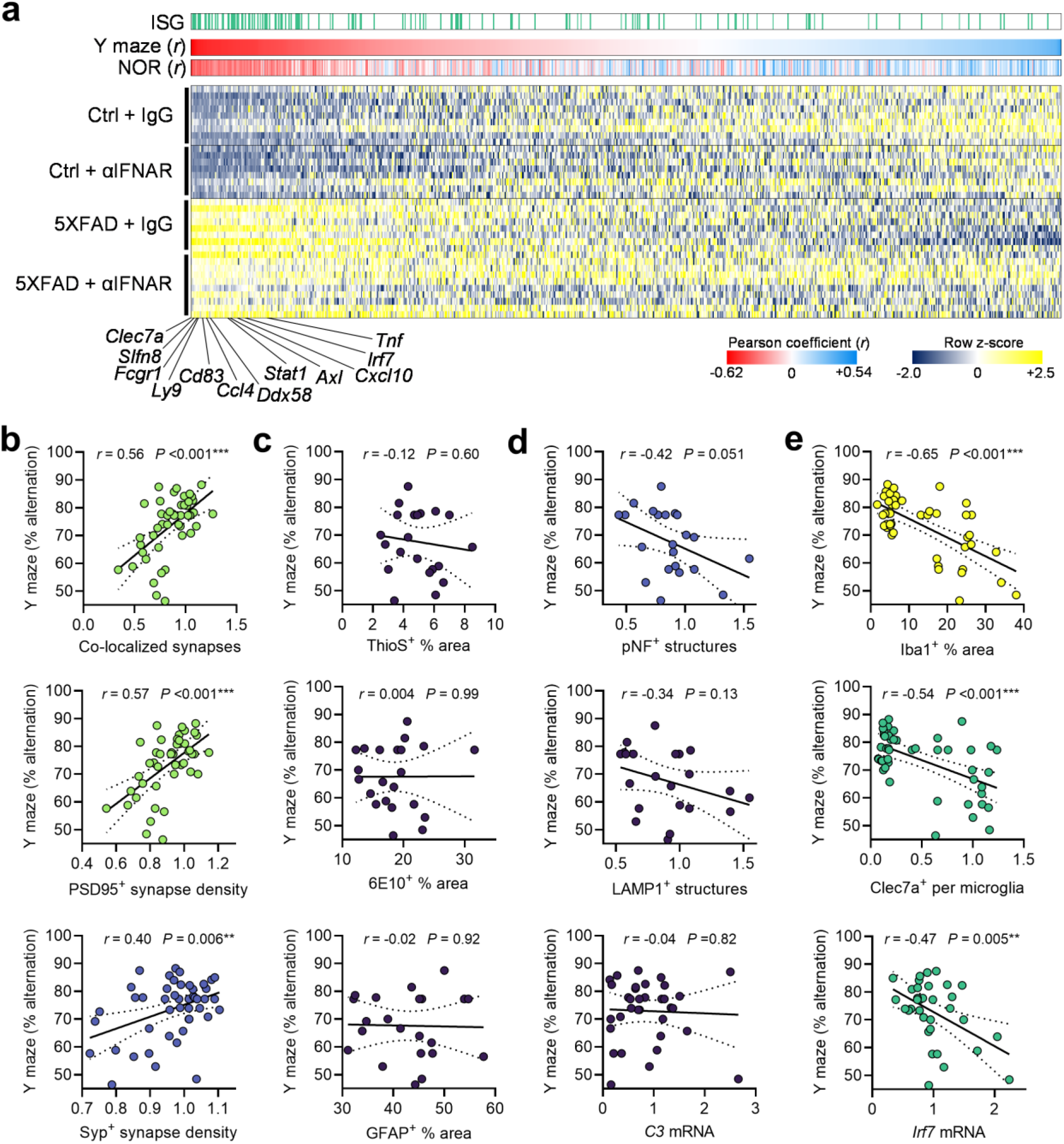
Memory impairment correlates with IFN signaling, synaptic pathology, and microglial reactivity. **a**, Nanostring gene expression analysis of hippocampal tissues from IgG- or αIFNAR-treated control or 5XFAD animals (1 animal per row; Ctrl + IgG, *n* = 9 animals; Ctrl + αIFNAR, *n* = 8 animals; 5XFAD + IgG, *n* = 8 animals; 5XFAD + αIFNAR, *n* = 10 animals). Top lanes: Genes identified as ISGs are marked with green. All genes are ranked (left to right) by their Pearson correlation (*r*) with impaired performance in Y maze. Correlations with NOR are also included. Several top inverse correlates are highlighted at bottom. **b-e**, Correlation analyses (Pearson *r*) of Y maze performance with numerous histopathological and transcriptional (*n* = 34 animals profiled) readouts from control and 5XFAD animals treated with IgG or αIFNAR (*n* = 46 total animals, groups as listed in **a**), including synaptic densities (**b**, *n* = 46 animals), plaque-related parameters (**c**, *n* = 22 animals), neuritic pathology parameters (**d**, *n* = 22 animals), and microglial activation markers (**e**, *n* = 46 animals). Simple linear regression lines (solid) with 95% CI intervals (dashed) were added to plots, as are Pearson *r* and associated *P* values. Plots were colored by *r* value as follows: 1 > *r* > 0.65, yellow; 0.65 > *r* > 0.55, light green; 0.55 > *r* > 0.45, dark green; 0.45 > *r* > 0.35, blue; *r* < 0.35, dark purple.

For neuropathology parameters, we found strong positive correlations of Y maze performance with PSD95/synaptophysin co-localized synapses, as well as with overall PSD95^+^ synapse density, and to a lesser extent with synaptophysin^+^ synapse density (Fig. 6b).

Although amyloid plaques represent a cardinal pathology of AD, we did not observe significant correlation of ThioS^+^ or 6E10^+^ plaque load with Y maze performance (Fig. 6c). Dystrophic axons, which were partially sensitive to IFN (Fig. 3e), displayed weak negative correlation with Y maze performance (Fig. 6d). Reactive astrocytes are known to participate in neurodegenerative processes. However, readouts of GFAP protein signal and *C3* mRNA, which is primarily produced by reactive astrocytes, did not correlate with memory capacity in this cohort (Fig. 6c,d).

Although microglial reactivity is highly influenced by β amyloidosis, whether activated states of these cells protect or harm the brain function remains controversial. We found that both overall microgliosis marked by Iba1 levels and DAM generation marked by Clec7a levels within microglia showed strong inverse correlations with Y maze performance (Fig 6e).

Hence, these findings suggest a potent negative impact of IFN and microglia on memory and cognition under the context of amyloid deposition.

### IFN signaling promotes microglia-mediated synaptic engulfment

Previously we showed that post-synaptic loss in 3-month-old 5XFAD brain was coupled with type I IFN-stimulated uptake by microglia (4). We did not detect any change in overall pre-synaptic density nor microglial pre-synaptic engulfment, implying that synapse loss is restricted to the post-synaptic compartment early in disease, and is IFN- and microglia-dependent. Given the concurrent synaptic deficits in pre- and post-synaptic elements (Fig. 2c) and seemingly differential cellular requirements for IFN signaling (Figs. 4, 5) in mid-stage 5XFAD mice, we sought to further investigate microglia in synapse modification.

First, synaptic engulfment assays showed that microglia in 5-month-old 5XFAD brain selectively engulfed enhanced amounts of PSD95^+^ puncta, an activity dependent on extracellular IFN (Fig. 7a) and necessarily mediated by IFN receptor in microglia (Fig. 7b). Consistent with this, more PSD95 was detected inside GFP^+^ microglia over GFP^-^ counterparts in plaque-rich regions of 5XFAD;MxG mice at 5 months (Fig. 7c). In contrast, no enhanced synaptophysin^+^ signal was detected inside microglia from 5XFAD brains, with or without IFN signaling, compared to control mice (Fig. 7a, b), suggesting a selective post-synaptic elimination by microglia, persisting at different stages of disease.

**Fig. 7:**
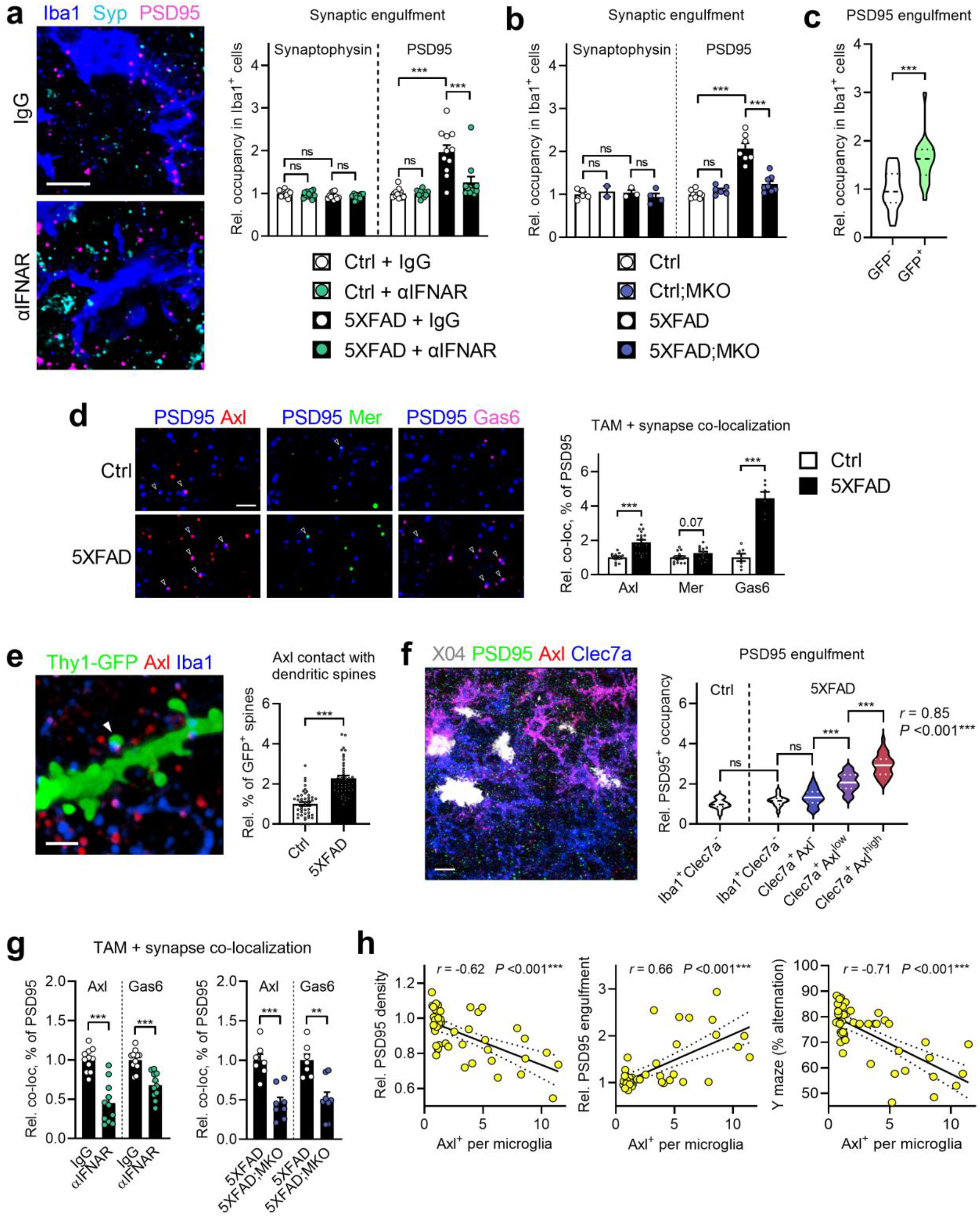
Post-synapses are preferential engulfed by IFN-stimulated Axl^+^ microglia. **a**, Representative images of microglia in relation to synaptic markers in subicula of treated 5XFAD animals as in Figure 2 (scale bar, 5 µm). Quantification of microglial engulfment of both markers. Ctrl + IgG, *n* = 13 animals; Ctrl + αIFNAR, *n* = 11 animals; 5XFAD + IgG, *n* = 11 animals; 5XFAD + αIFNAR, *n* = 11 animals. Data represent means and s.e.m. Statistics were performed with ordinary one-way ANOVA (Syp: *P* = 0.0717, F_42_ = 2.510; PSD95: *P* < 0.001, F_42_ = 19.23) and Bonferroni’s multiple-comparisons test. ns, not significant; ^***^*P* < 0.001. **b**, Quantification of microglial engulfment of pre- and post-synaptic puncta in subicula of Ctrl (*n* = 5 animals for Syp, *n* = 9 animals for PSD95), Ctrl;MKO (*n* = 2 animals for Syp, *n* = 6 animals for PSD95), 5XFAD (*n* = 3 animals for Syp, *n* = 7 animals for PSD95), and 5XFAD;MKO (*n* = 4 animals for Syp, *n* = 8 animals for PSD95). Data represent means and s.e.m. Statistics were performed with ordinary one-way ANOVA (Syp: *P* = 0.688, F_10_ = 0.5038; PSD95: *P* <0.001, F_26_ = 44.28) and Bonferroni’s multiple-comparisons test. ns, not significant; ^***^*P* < 0.001. **c**, Quantification of relative uptake of PSD95^+^ post-synaptic puncta by GFP^-^ (*n* = 19 cells) and GFP^+^ microglia (*n* = 21 cells) from 5-month-old 5XFAD;MxG animals (*n* = 3 animals). Data are presented as a violin plot with medians (dashed lines) and quartiles (dotted lines). Statistics were performed with two-tailed *t*-test. ^***^*P* < 0.001. **d**, Representative high-magnification confocal images of PSD95^+^ post-synapses in proximity (≤200 nm) to TAM receptors Axl and Mer, and TAM ligand Gas6 (scale bar, 2 µm). Quantification of co-localization, as relative percent of PSD95^+^ puncta, of the three molecules with PSD95 (Axl + PSD95: *n* = 13 Ctrl images, *n* = 17 5XFAD images; Mer + PSD95: *n* = 14 Ctrl images, *n* = 15 5XFAD images; Gas6 + PSD95: *n* = 7 Ctrl images, *n* = 6 5XFAD images). Data represent means and s.e.m. of images from *n* = 4 animals per genotype. Statistics were performed with two-tailed *t*-tests. ^***^*P* < 0.001. **e**, Representative high-magnification confocal image of Axl^+^ microglial processes contacting dendritic spines (arrow) in a 5XFAD mouse on the Thy1-eGFP background (scale bar, 2 µm), and quantification of relative frequency of observed contacts between control (*n* = 42 dendrites >10 µm long from *n* = 8 animals) and 5XFAD mice (*n* = 44 dendrites >10 µm long from *n* = 8 animals). Data represent means and s.e.m. Statistics were performed with two-tailed *t*-test. ^*^*P* < 0.001. **f**, Representative confocal image of Clec7a^+^ microglia with varying degrees of Axl expression in relation to PSD95^+^ post-synapses in a 5-month-old 5XFAD animal (scale bar, 20 µm). Histological analysis of single microglia in both control and 5XFAD brains stratified by levels of Clec7a and Axl expression, showing relative amounts of PSD95^+^ synaptic uptake in each category. Ctrl Iba1^+^Clec7a^-^, *n* = 17 cells; 5XFAD Iba1^+^Clec7a^-^, *n* = 19 cells; 5XFAD Clec7a^+^Axl^-^, *n* = 37 cells; 5XFAD Clec7a^+^Axl^low^, *n* = 35 cells; 5XFAD Clec7a^+^Axl^high^, *n* = 32 cells; all cells combined from *n* = 2 Ctrl animals and *n* = 3 5XFAD animals at 5 months. Data are presented as a violin plot with medians (dashed lines) and quartiles (dotted lines). Statistics were performed with ordinary one- way ANOVA (*P* <0.001, F_135_ = 95.97) with Bonferroni’s multiple-comparisons test. ns, not significant; ^***^*P* <0.001. Pearson *r* at right was calculated by correlation analysis of Axl expression and PSD95^+^ uptake in all Clec7a^+^ cells. **g**, Quantified co-localization of Axl and Gas6 with PSD95, as relative percent of the synaptic puncta, in 5XFAD animals treated with IgG (*n* = 11 animals) or αIFNAR (*n* = 11 animals), and in *Ifnar1* microglia conditional KO lines (5XFAD, *n* = 7 animals; 5XFAD;MKO, *n* = 8 animals). Data represent means and s.e.m. Statistics were performed with two-tailed *t*-tests. ^**^*P* < 0.01; ^***^*P* < 0.001. **h**, Correlation analyses (Pearson *r*) of post-synaptic density, engulfment, and Y maze performance with the extent of Axl expression in microglia by histological analysis in 5XFAD animals treated with IgG or αIFNAR (*n* = 46 total animals, groups as listed in Figure 6a). Simple linear regression lines (solid) with 95% CI intervals (dashed) were added to plots, as are Pearson *r* and associated *P* values.

We previously showed that IFN-activated microglia rapidly remove dendritic spines in a complement C3-dependent manner (4). Although IFN was sufficient in inducing many members of the complement cascade in wild-type mice, blockade of extracellular IFN or genetic IFN receptor ablation in 5XFAD mice did not reduce complement transcription (Fig. S7a,b), consistent with unchanged C3 protein in astrocytes (Fig. S3b). This indicates that signals other than IFN may play a role in chronic complement activation in older 5XFAD mice.

Perineuronal nets (PNN) are extracellular matrix structures that enwrap and stabilize neuronal synapses, loss of which in 5XFAD was shown to be mediated by microglia (17). Although *Wisteria floribunda* agglutinin (WFA) staining confirmed a significant reduction of PNN structures in disease, IFN blockade did not appear to affect their levels (Fig. S7c), excluding a direct link between IFN and PNN modification.

Axl is a member of the TAM (Tyro3, Axl, and Mer) family RTKs that play important roles in phagocytosis of apoptotic cells (18). Recently, a plaque-centric expression pattern of TAM receptors and their ligand Gas6 was reported to engage microglia with amyloid plaques in a largely Mer-dependent manner (19). Given the high sensitivity of microglial Axl to IFN signaling, we investigated its relation to synapses, together with Mer and Gas6. Employing high-magnification confocal imaging, we detected specific, punctate signals for Gas6 as well as both Axl and Mer in wild-type brain (Fig. S7d), which interestingly displayed non-random co-localization with synaptic puncta (Fig. S7e), indicating a physiological interaction of TAM molecules with synapses. In diseased brain, we observed notable Gas6 deposition on amyloid plaques and enhanced Mer expression in plaque-associated microglia (Fig. S7f,g), in agreement with Huang et al (19). However, unlike Axl, Mer expression in microglia, as well as extent of Gas6 deposition on plaques, were not IFN-dependent. At the synaptic structures, we found substantial Gas6 deposition on PSD95^+^ synaptic terminals in 5XFAD brains, which was accompanied by significantly increased Axl, but not Mer, co-localization with PSD95 (Fig. 7d). To visualize the physical relationship between Axl and synapses, we analyzed dendritic spines of 5XFAD mice containing the Thy1-eGFP reporter. High-magnification confocal imaging revealed the formation of contact points between GFP^+^ dendritic spines and Axl^+^ microglial processes, which were significantly more frequent along dendrites in 5XFAD mice (Fig. 7e), substantiating a direct contact of Axl with synapses.

To explore the role of Axl receptor in synapse uptake, we examined PSD95 engulfment by different subpopulations of microglia from control and 5XFAD brains, particularly Clec7a^+^ plaque-associated microglia with varying expression of Axl, and detected robust per-cell correlation of Axl and PSD95 occupancy in microglia (Fig. 7f). To test whether Axl and/or Gas6 localization to synapses is dependent on IFN signaling, we measured the frequency of Axl/PSD95 and Gas6/PSD95 co-localization in 5XFAD mice treated with IgG or αIFNAR, and found that colocalization of both Axl receptor and Gas6 ligand to synapses was reduced with IFN blockade, a finding which was mirrored by microglial conditional *Ifnar1* deletion (Fig. 7g), suggesting a reversible, IFN-induced post-synaptic engulfment machinery in microglia during disease. In line with this, Axl protein and mRNA levels displayed a strong positive correlation with PSD95 engulfment, and negative correlations with PSD95 density and Y maze performance (Fig. 7h, S6a). Altogether, these data pinpoint an IFN-instructed synapse elimination program in microglia that compromises memory.

## Discussion

Beyond antiviral function, type I IFN is linked to cognitive and neuropsychiatric dysfunction in various clinical contexts (4, 20). Previous studies describe that IFN modifies the brain through microglia activation, neural stem cell dysfunction, and disruption of whole-brain functional network connectivity (4, 21). Of the neuropathological features of AD, synapse loss appears early and correlates most strongly with dementia, and thus represents a key step of the disease process (22). Our current study reveals for the first time discrete and coordinated actions of IFN-stimulated brain cells in compromising synapses, the central cause of memory impairment, under the sterile inflammatory condition initiated by AD pathology (Fig S8).

A growing number of microglia populations are being identified by scRNA-seq analyses, revealing different activation states under various physiological or pathological conditions (23). While cells enriched with DAM markers were identified first, IRMs were subsequently recognized as a distinct subset of microglia arising in AD and brain aging (6, 7, 24). Moreover, a microglial proteome analysis revealed that IFN pathway was activated early and persisted in murine Aβ models (25), in line with our findings that microglial IFN response universally accompanies brain amyloidosis *in vivo* (4). Using a genetically-encoded IFN-responsive reporter system, we documented an age-dependent, brain-wide, and profound accrual of brain cells responding to IFN signaling activation in the 5XFAD model. In young mice, a sparse population of microglia were the principal IFN-responsive cell-type, consistent with the results obtained with Stat1 staining (4). By 5 months, despite overwhelming presence of NA^+^ plaques, no more than half of plaque-associated Clec7a^+^ microglia expressed GFP, revealing an interesting aspect of microglial heterogeneity. Given the higher percentage of cells turning green at older age, microglia seem to be activated by IFN continuously as amyloidosis progresses. It is also worth noting that over 90% of GFP^+^ microglia retained Clec7a expression, which implies that microglia maintain DAM markers after IFN activation. As GFP^+^ microglia accumulated alongside the increasing plaque load, other brain cells also became GFP^+^, revealing a more complex IFN response than previously appreciated. Many brain cell types participate in plaque formation and neuritic pathology, a process marked by a multicellular co-expression network of plaque-induced genes (PIGs) (26). We found that 22 of the 57 PIG module are CNS ISGs (4, 27, 28), many of which overlap with the markers of DAM and neurotoxic reactive astrocytes (29) (Fig. S9), highlighting a profound influence of IFN in the dysregulated cellular network in the vicinity of plaques.

We obtained apparently conflicting results on whether IFN signaling affects plaque pathology: blocking IFN receptor did not (Fig. 2d), while neural *Ifnar1* deletion partially reduced plaque load (Fig. 5b). The latter observation was correlated with significantly tempered Ifitm3 expression, in keeping with the activity of Ifitm3 in promoting APP cleavage and amyloid pathology (5). Paradoxically, Ifitm3 was unaltered with IFN blockade (Fig. S2d), implying a difference between extracellular IFN blocking and genetic ablation. One possible explanation comes from clinical observation with therapeutic αIFNAR antibodies that IFN has more persistent effects in cells devoid of negative IFN regulators ISG15 and USP18 (30). Interestingly, cortical neurons do not express Isg15 or Usp18 (both are ISGs) even after IFN exposure, a contrast to microglia (27, 28). Given the sensitive neuronal response of Ifitm3 induction by IFN (Fig. S5c), it is plausible that, under IFNAR blockade, residual IFN receptor signaling was sufficient to maintain the levels of Ifitm3. Of note, not all neuronal IFN signaling escaped extracellular blockade, as neuronal pStat1 and total Stat1 proteins were similarly reduced by antibody-mediated blockade (Figs. S2e, 3a) and genetic ablation of *Ifnar1* in neural cells (Fig. 5a,e).

Another unique pathological hallmark of AD is the swollen pre-synaptic dystrophic neurites surrounding amyloid plaques, which accumulate APP as well β- and γ -secretases, and serve as localized sites of Aβ generation and release (31, 32). As we have shown (Fig. 5), neural IFN signaling was required for Ifitm3 expression, which is known to enhance γ -secretase activity. Remarkably, β-secretase expression is also reportedly regulated by interferon and Stat1 (33, 34), implying a sweeping effect of IFN on APP processing. Overall, our results support a feed-forward Aβ-plaque-IFN-Aβ loop whereby inflammation stimulates factors that exacerbate AD pathology.

We discovered that synaptophysin^+^ boutons were selectively diminished by IFN signaling in neural-derived cells (Fig. 5d), in sharp contrast to the regulation of post-synaptic densities (Fig. 4d). A Jak2-Stat1 axis was recently identified as a major neuron-autonomous determinant to eliminate inactive synapses *in vivo* (16). Interestingly, Stat1 functions not only as a negative regulator of spatial memory formation in wild-type mice, but is also a key mediator of Aβ-induced learning and memory deficits (35, 36). We found increased pre-synaptic pStat1 in 5XFAD, which was reduced upon IFN blockade or ablation (Figs. 5e, S2e). Collectively, the experimental evidence strongly supports a novel neuronal IFN-Stat1 axis that pathogenically modulates the pre-synapse in AD (Fig S8).

While microglia prune synapses during normal CNS development, excessive removal can result in pathological synapse loss in diverse neurological and neuropsychiatric diseases (37). In β-amyloidosis models, germline C3 deficiency protects from loss of synapses and neurons (38, 39), and microglia engulf C1q-tagged post-synaptic components early in the disease (40). Despite the strong relationship between IFN and complement in young 5XFAD mice, we found unexpectedly that IFN signaling became dispensable in eliciting complement expression in 5- month-old animals, likely reflecting the influences of other prevailing proinflammatory signals. It should be noted that, in the same cohort, IFN blockade was effective to blunt *Tnf* and *Clec7a* expression, similar to the treatment effects in 10-to 12-month-old APP^NL-G-F^ mice (4).

Besides complement, microglia use myriad other surface receptors to engulf or otherwise limit synapses (41-44). Interestingly, several synapse-eliminating receptors recognize a common neuronal cue (45): phosphatidylserine (PS), a well-known “eat-me” signal for phagocytosis. The principal myeloid phagocytic receptors, Axl and Mer, detect PS exposed on apoptotic cells via their ligand Gas6, which displays high affinity towards PS (18). In CNS, Mer facilitates astrocytic phagocytosis of synapses in developing and adult brain (46), and can also engage plaques in AD (19). Yet, the function of Axl in brain, despite its prominent upregulation in plaque-associated microglia in AD, is unknown. Intriguingly, we discovered a highly IFN-dependent Axl expression in microglia surrounding amyloid plaques (Fig 3,4), As reported (19), we found *Mertk* mRNA positively correlated with dense-core plaques (Fig. S6c). Contrary to Mer, IFN blockade and microglial *Ifnar1* deletion effectively reduced Axl levels but failed to modify the plaques, in line with *Axl* deficiency in APP/PS1 model (19). On the other hand, we demonstrate for the first time direct contact between Axl and synapses, which were highly tagged by Gas6 in β-amyloidosis, and enrichment of synaptic material inside Axl^+^ microglia, all of which were IFN-dependent. While Mer expression displayed no association with either memory or synapse levels, Axl abundance was robustly and inversely correlated with memory performance (Fig. S6a,b). Interestingly, both soluble AXL and GAS6 levels increase in cerebrospinal fluid of AD patients (47, 48), consistent with the elevated Axl expression in human AD (4)(Fig. S7h). These intriguing findings warrant further characterization of a pathogenic involvement of microglial Axl in AD.

We present evidence that IFN signaling plays a role in phosphorylation of endogenous tau at neuritic plaques (Fig. 3), which constitutes a major type of AD-relevant tau pathology and, notably, has been shown to enable AD-tau spreading *in vivo* (49). Although uncertain how IFN modulates tau together with other peri-plaque dystrophies, we were intrigued by the report that increased IRMs were associated with heightened endogenous tau phosphorylation in *Trem2*-deficient APP/tau double transgenic mice (50). Hence, the mechanism by which IFN-mediated signaling connects amyloid and tau pathologies in AD, and whether IFN pathway represents a feasible therapeutic target, are of great interest.

## Acknowledgements

The study was funded by NIH grants AG057587 (W.C. and H.Z.); AG020670 (H.Z.), AG062257 (H.Z.), NS093652 (H.Z.), BrightFocus ADR A20183775 (W.C.) and Brown Foundation 2020 Healthy Aging Initiative (W.C.). We acknowledge technical assistance from Haiying Liu, Nadia Aithmitti, and Bianca Contreras, and express gratitude to Dr. Andre Catic for generously providing MxG founder mice.

## Author Contributions

Conceptualization, E.R.R. and W.C.; Methodology, E.R.R., G.C. and S. L.; Formal Analysis, E.R.R..; Investigation, E.R.R., G.C. and S. L.; Resources, N.E.P.; Writing-Original Draft, E.R.R., and W.C.; Writing-Review & Editing, E.R.R. and W.C.; Visualization, E.R.R.; Supervision, W.C.; Project Administration, W.C.; Funding Acquisition, H.Z., and W.C.

